# New insights into the role of oxytocin signaling in preeclampsia

**DOI:** 10.1101/2025.04.10.648289

**Authors:** Sung-Min An, Jea Sic Jeong, Min Jae Kim, Da Som Kim, So Young Kim, Hyeon-Gu Kang, Beum-Soo An, Seung-Chul Kim

## Abstract

**Background:** Oxytocin signaling may play an important role in the human placenta during pregnancy, where abnormal function can cause pregnancy complications such as preeclampsia, a hypertension syndrome with elusive etiology, treatment, and diagnosis. As pregnancy progresses, the clinical effects of high blood pressure tend to worsen. Therefore, an early diagnosis of preeclampsia is necessary for a successful pregnancy.

**Method:** This study examined placenta and plasma samples from normal and preeclamptic pregnancies, approved by the Institutional Review Board. qRT-PCR, western blot, ELISA, gelatin zymography, and ChIP were performed on placental tissues. JEG-3 cells and HUVECs were cultured under normoxia or hypoxia with oxytocin or atosiban. siRNA/miRNA transfections and RUPP rat models assessed oxytocin’s effects. Data were analyzed using ANOVA and Duncan’s test (P ≤ 0.05) in SPSS.

**Result:** Here, we assessed the effects of oxytocin signaling on key placental functions and its correlation with preeclampsia, revealing a significant association with placental dysfunction. Our findings revealed that oxytocin increased trophoblast invasion via ERK1/2- and JNK/AP-1-dependent pathways. Notably, oxytocin signaling was inhibited by hsa-miR-193b-5p and hypoxia. Administering atosiban to pregnant rats hindered placental invasion activity, mirroring symptoms of preeclampsia, and disrupted placental development. Finally, we examined oxytocin signaling in patients with preeclampsia by analyzing oxytocin levels in the first trimester. Oxytocin levels were higher in those who later developed preeclampsia than normal pregnant women.

**Conclusion:** These results indicate that changes in oxytocin signaling are related with preeclampsia throughout pregnancy, emphasizing its potential as a novel early diagnostic biomarker.

## Background

Preeclampsia (PE) is a multisystem disorder that is characterized by the new onset of hypertension during pregnancy, including elevated blood pressure and proteinuria. It affects approximately 2%–8% of pregnancies and typically develops after 20 weeks of gestation ^1^. Clinical features of PE include hypertension (systolic blood pressure ≥140 mmHg or diastolic blood pressure ≥90 mmHg), proteinuria (≥300 mg/24 h), placental hypoxia, endothelial dysfunction, end-organ ischemia, and increased vascular permeability. These clinical manifestations can substantially contribute to maternal and fetal morbidity and mortality ^2^. Several theories have been proposed to explain the etiology and pathogenesis of PE; however, none have been able to satisfactorily account for the numerous risk factors associated with this condition ^3^.

Several studies have reported that impaired trophoblast cell invasion of the uterine blood vessels is a hallmark of PE ^4,5^. In a normal pregnancy, proper trophoblast invasion leads to the formation of spiral arteries, which facilitate gaseous exchange and create a placental layer characterized by high flow and low resistance ^6^. However, in cases of PE, abnormal trophoblast invasion can occur early during pregnancy, resulting in abnormal placental formation. Placentas from patients with PE often exhibit reduced trophoblast invasion at the decidua and myometrium levels, as well as an abnormal state of trophoblast cells in the generation of spiral arteries ^7^.

The infiltration of the trophoblast into the decidual stroma is facilitated through the degradation of the extracellular matrix (ECM) proteins via the action of proteolytic enzymes. Matrix metalloproteinases (MMPs) are the primary regulators of trophoblast invasion ^8^. Specifically, MMP-2 (gelatinase A) and MMP-9 (gelatinase B) play essential roles in endometrial tissue remodeling and are abundantly expressed in the invading extravillous trophoblast cells, which contributes to their invasiveness ^9–11^. MMPs have been implicated in embryo implantation during the early stages of pregnancy and childbirth, as well as successful pregnancy maintenance ^12^. Therefore, a comprehensive investigation of placental penetration ability, which is considered to be a significant factor correlated with PE, is crucial for understanding the etiology of PE and the establishment of innovative treatment strategies ^13^. Because the pathogenesis of PE remains under investigation and the only known treatment is delivery, it is essential to identify and prevent PE early during pregnancy by developing novel PE biomarkers. Currently, there are no definitive biomarkers available for the early diagnosis of PE, thus emphasizing the need to identify promising new biomarkers for this disease ^14^.

Oxytocin (OXT), a nonapeptide hormone, was initially discovered as a neurohypophyseal hormone ^15^. It is primarily synthesized by magnocellular neurons that are located in the supraoptic nucleus and paraventricular nucleus of the hypothalamus, with their axons terminating in the posterior lobe of the pituitary gland ^16^. In addition, OXT is synthesized by various peripheral tissues, including the uterus, amnion, corpus luteum, gonads, gut, heart, and placenta ^16^. Following synthesis and release, OXT binds to the OXT receptor (OXTR) and plays various important roles in physiological processes, particularly in the reproduction process, such as maternal behavior, labor, and lactation ^17^.

In our previous study, we demonstrated that both OXT and OXTR are highly expressed in the human placenta, and OXT was distributed in both syncytiotrophoblasts and cytotrophoblasts ^18^. Furthermore, OXT levels were found to progressively increase in the maternal blood throughout the pregnancy, and recent research has identified a gradual rise in plasma OXT levels with advancing gestational weeks ^16,18^. Based on these findings, we hypothesize that OXT plays a crucial role in the placenta for successful pregnancy progression. Given that PE is a complex condition that results from multiple concurrently occurring mechanisms, we propose that the impairment of placental OXT signaling could contribute to the underlying pathogenesis of the various clinical symptoms of PE.

In the present study, we investigated the correlation between OXT signaling and PE, a condition that is closely associated with placental dysfunction, at both the *in vitro* and *in vivo* levels. To the best of our knowledge, no previous study has compared plasma OXT concentrations between healthy pregnant women and those with PE. Therefore, the objective of this study was to assess the potential use of OXT as a biomarker for the early diagnosis of PE by measuring maternal plasma OXT concentrations during the early stages of pregnancy.

## Methods

### Human placenta tissue and blood collection and processing

This study was approved by the Institutional Review Board (IRB) of the Pusan National University Hospital Clinical Trial Center (1302-005-015), and all participants provided written informed consent. Human placenta tissues and plasma samples were collected and immediately stored at −80°C until evaluation. The samples were divided into groups of normal women (n = 21) with no preexisting clinical conditions such as diabetes, hypertension, or autoimmune disease and patients with PE (n = 20). The samples were obtained between 29 and 40 weeks of gestation and were provided by the Biobank of Pusan National University Hospital (Busan, Korea), a member of the Korea Biobank Network. Patients with PE were defined as having systolic and diastolic blood pressures ≥140 and ≥90 mm Hg, respectively, which were measured at least 6 h apart, in addition to proteinuria ≥300 mg/24 h or ≥1+ as determined by the dipstick method. The clinical characteristics of the patients are presented in Table S1.

In addition, plasma samples from the first trimester of pregnancy were collected and stored from normal women (n = 16) and patients with PE (n = 40) with IRB approval (2006-030-092). Plasma samples were obtained between 12 and 13 weeks of gestation, and the clinical characteristics of the patients are presented in Table S2.

### Cell preparation and culture conditions

The JEG-3 human choriocarcinoma-derived cell line (Korean Cell Line Bank, Seoul, Republic of Korea) was cultured in DMEM. HUVECs were grown in the Endothelial Cell Growth Medium 2 Kit (Promo Cell, Heidelberg, Germany, Cat. no. C-22211) in culture flasks. All growth media contained 10% FBS and 1% streptomycin and penicillin (Welgene, Inc.). Both cells were cultivated at 37°C in a humidified atmosphere maintained with 5% CO_2_.

### Animal experiments

Animal studies were approved (approval number: PNU-2022-0161) by the Ethics Committee of Pusan National University (Busan, Republic of Korea). Pregnant female Sprague–Dawley (GD 1) rats (n = 35) were purchased from Samtako (Osan, Republic of Korea) twice in accordance with the experimental schedule. All rats were housed in standard laboratory conditions with controlled temperature and humidity, a 12 h/12 h light/dark cycle, and free access to food and water at the Pusan National University Laboratory Animal Resources Center. This center is accredited by the Korea Food and Drug Administration in accordance with the Laboratory Animal Act (Accredited Unit Number: 000231) and the Association for Assessment and Accreditation of Laboratory Animal Care International according to the National Institutes of Health guidelines (Accredited Unit Number: 001525). After acclimatization, in the first animal experiment, rats were divided into six groups as follows: control group (n = 5), atosiban low-dose group (0.6 mg/kg/day; n = 5), atosiban high-dose group (1.2 mg/kg/day; n = 5), SHAM group (n = 5), and RUPP group (n = 5). Atosiban groups were administered atosiban by subcutaneous (SC) injection daily from GD 10 to 17 ^19^.

For the second animal experiment, to further evaluate the effect of atosiban during early pregnancy, the rats were divided into two groups: the control group (n = 5) and the atosiban high-dose group (1.2 mg/kg/day; n = 5). Atosiban was SC injected daily at GD 3 to 7. In all experiments, the control group was administered 0.9% saline. The dosage was adjusted according to changes in maternal BW. At the end of the experiments, the rats were euthanized in a gradually filled CO_2_ gas chamber with a flow rate of ≤30% CO_2_ of the chamber volume/min. BW, clinical signs, and abnormal behaviors were recorded daily throughout the experimental period.

### Data analysis

SPSS for Windows version 20.0 (SPSS Inc., Chicago, IL, USA) was used for statistical analysis. All experiments were performed in triplicate, and the results are presented as the mean ± standard deviation (SD). The data were analyzed using one-way analysis of variance, and differences between the mean values were separated using Duncan’s test at a significance level of P ≤0.05.

## Results

### OXT signaling is significantly associated with trophoblast cell permeability

The invasion of placental cells is a crucial process for the successful development and maintenance of pregnancy ^21^. To investigate the role of OXT signaling in trophoblast invasion, we performed a transwell invasion assay using JEG-3 cells. As depicted in Figure 1A, OXT significantly increased cell invasion across the Matrigel-coated membrane. Moreover, OXT treatment resulted in a significant increase in the mRNA and protein expression levels of MMP-2 and MMP-9 in JEG-3 cells (Figure 1B and C).

**Figure 1.**
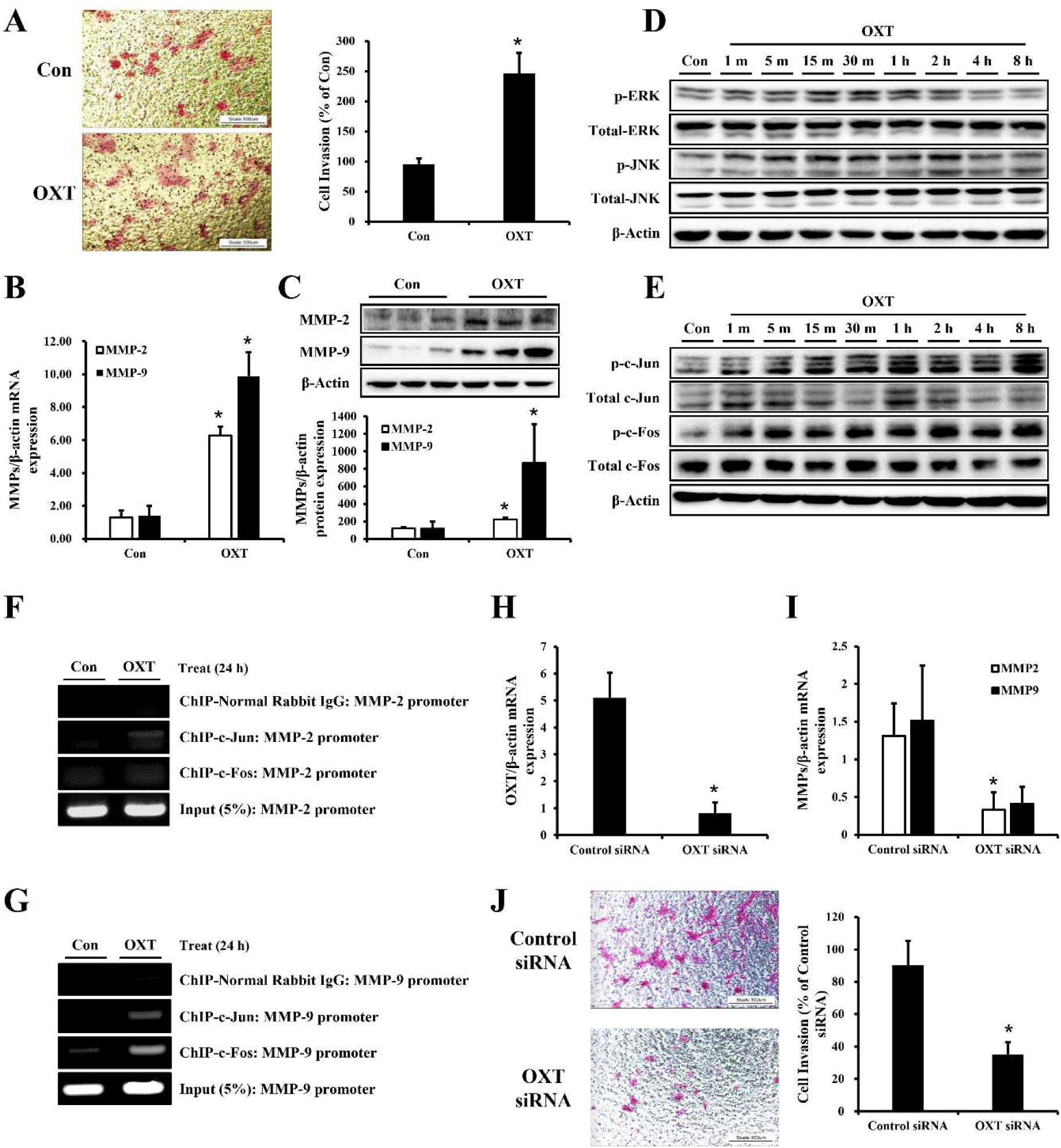
Effects of oxytocin (OXT) on the permeability of trophoblast cells. (A) JEG-3 cells were treated with OXT, and a transwell invasion assay was performed to analyze the invasive ability of the JEG-3 cells after OXT treatment. Representative images of the filters contain invaded cells from Matrigel invasion assays. The invasion ratio of the cells was measured. Scale bar = 100 µm. The data are expressed as the mean ± standard deviation (SD), *P ≤ 0.05, relative to the control group. (B) Total mRNA was extracted from the OXT-treated JEG-3 cells. MMP-2 and MMP-9 transcription levels were examined by qRT-PCR. (C) Western blot analysis was performed to analyze the protein expression levels of MMP-2 and MMP-9 after OXT treatment of JEG-3 cells, and the values are represented as schematic graphs. The mRNA and protein expression levels were normalized to those of β-Actin. The data are expressed as the mean ± SD, *P ≤ 0.05, relative to the control group. (D and E) Time course of western blot analysis. JEG-3 cells were treated with OXT for the indicated times. Cell lysates were sequentially immunoblotted with indicated antibodies to detect the phosphorylated and total proteins of ERK1/2, JNK, c-Jun, and c-Fos. β-Actin was used as the loading control. (F and G) The association of the AP-1 site (c-Jun/c-Fos) with the human MMP-2 and MMP-9 promoters was analyzed by a chromatic immunoprecipitation (CHIP) assay in the JEG-3 trophoblast cells using antibodies against c-Jun and c-Fos or an unrelated IgG antibody. Amplified MMP-2 (F) and MMP-9 (G) promoter fragments of the AP-1-element-containing sequence are shown. The amount of DNA in the input confirms the equal loading of chromatin. (H) The siRNA-mediated knockdown of OXT was performed as described in the Materials and Methods section. After transfection, qRT-PCR was performed to analyze the knockdown efficiency. The data are expressed as the mean ± SD, *P ≤ 0.05, relative to the non-targeting control siRNA group. (I) Following the siRNA-mediated knockdown of OXT, qRT-PCR was performed to analyze the gene expression of MMP-2 and MMP-9. β-Actin was used to normalize the mRNA expression. The data are expressed as the mean ± SD, *P ≤ 0.05, relative to the non-targeting control siRNA group. (J) A transwell invasion assay was performed to analyze the invasive ability of the JEG-3 cells after control or OXT siRNA transfection. The invasion ratio of the cells was measured. Scale bar = 100 µm. The data are expressed as the mean ± SD, *P ≤ 0.05, relative to the nontargeting control siRNA group.

To further elucidate the signaling events underlying MMP gene regulation, we examined the effects of OXT on AP-1, a key transcription factor for MMP expression ^22^. Previous studies have highlighted the essential roles of the ERK1/2 and JNK pathways in MMP expression through AP-1 ^23,24^. As shown in Figure 1D, OXT significantly enhanced the phosphorylation of ERK1/2 and JNK in JEG-3 cells. The activation of phosphorylation by OXT treatment occurred rapidly, within 15 min, and was sustained for several hours. Furthermore, OXT treatment increased the phosphorylation of c-Jun and c-Fos, which are components of the AP-1 pathway, in a time-dependent manner (Figure 1E). To confirm the involvement of AP-1 in the OXT-induced MMP activation, we performed a chromatic immunoprecipitation (CHIP) assay to examine the recruitment of c-Jun and c-Fos to the MMP promoter region. OXT stimulated the binding of c-Jun to the MMP-2 promoter, whereas both c-Jun and c-Fos exhibited increased binding to the MMP-9 promoter (Figure 1F and G). These findings suggest that the OXT-induced activation of the ERK1/2, JNK, and AP-1 signaling pathways leads to the increased expression of MMP, thereby promoting the invasion of JEG-3 cells.

In our previous study, we demonstrated that OXT is evenly produced in human placental trophoblast cells and plays a critical endogenous role during pregnancy ^18^. To elucidate the endogenous role of OXT in the permeability of trophoblasts, we transfected JEG-3 cells with OXT siRNA. The efficacy of siRNA targeting OXT was demonstrated by an 80% reduction in OXT gene expression (Figure 1H). In addition, the downregulation of OXT significantly suppressed mRNA expression of MMP-2 but not MMP-9 (Figure 1I), and the invasion activity of JEG-3 cells was significantly reduced upon OXT knockdown (Figure 1J).

These results suggest that endogenous OXT plays a significant role in the penetration ability of human trophoblast cells, and the absence of OXT in placental cells can result in a significant decrease in MMP gene regulation, which is closely associated with the invasion process that is essential for placental development during pregnancy.

### PE-associated placenta-derived hsa-miR-193b-5p decreases trophoblast cell permeability by reducing OXT signaling

Inadequate trophoblast penetration has been associated with pregnancy-specific disorders such as PE ^25^. Previous studies have suggested that the abnormal expression of placental-derived miRNAs contributes to the development of PE ^26,27^. In this study, we used an online database for miRNA target prediction and selected hsa-miR-193b-5p as a candidate miRNA from among the PE-associated placental-derived miRNAs that could affect OXT signaling.

To investigate the influence of miR-193b-5p on the expression of OXT in JEG-3 cells, we transfected the cells with an hsa-miR-193b mimic and assessed the transfection efficiency using qRT-PCR. Our data demonstrated the successful expression of hsa-miR-193b after transfection compared with the negative control group (Figure 2A), and its overexpression resulted in the downregulation of OXT (Figure 2B). However, hsa-miR-193b did not alter the level of OXT released from JEG-3 cells or the mRNA and protein levels of OXTR (Figure S1A, Figure 2C and D).

**Figure 2.**
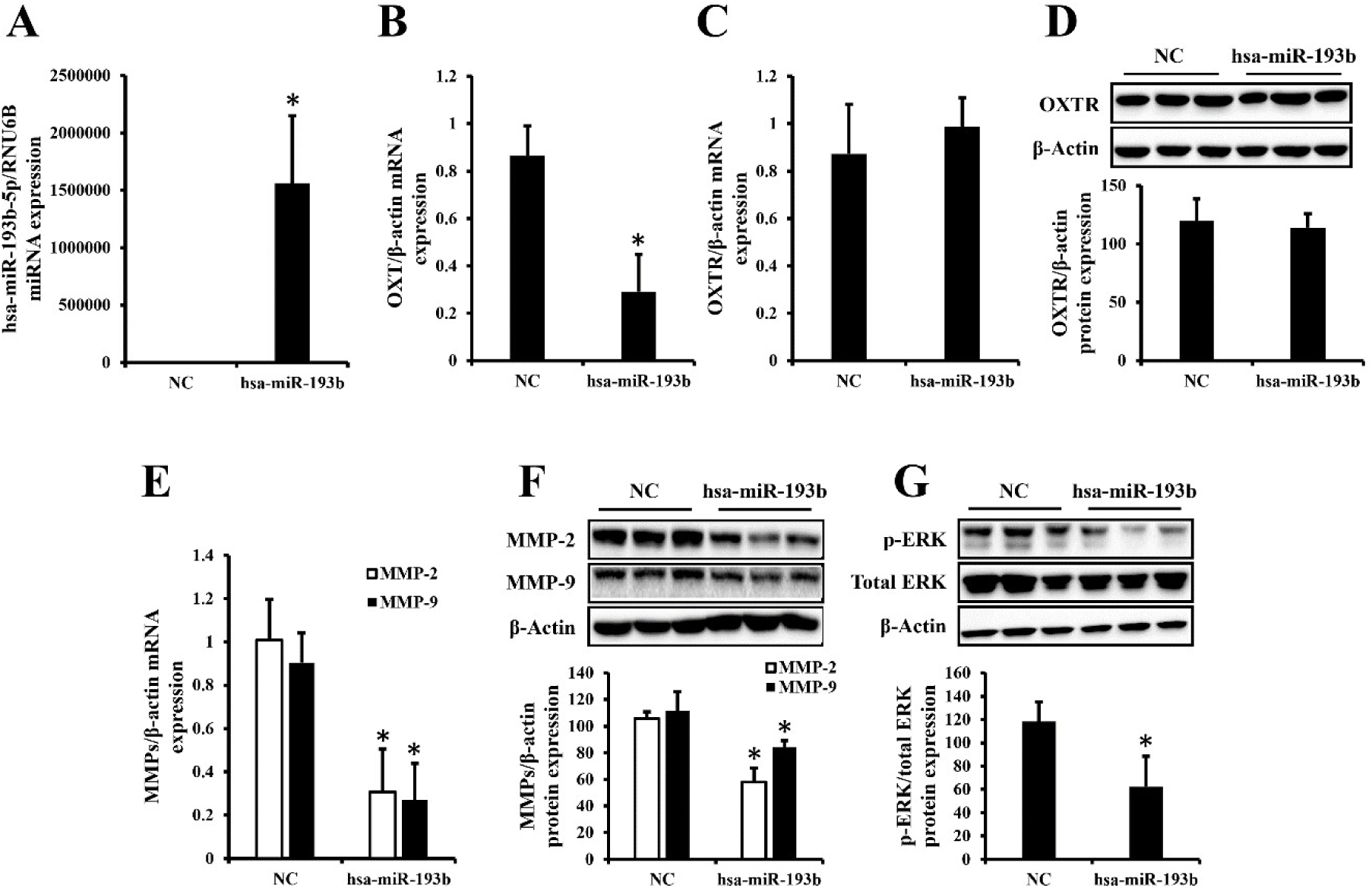
Overexpression of hsa-miR-193b-5p in trophoblast cells decreases invasion ability by inhibiting oxytocin (OXT) signaling. (A) The JEG-3 cells were transfected with either a negative control or a hsa-miR-193b mimic. After transfection, qRT-PCR was performed to evaluate the transfection efficiency of hsa-miR-193b-5p using the small nuclear RNA RNU6B levels as the reference. (B-D) Transcriptional and translational levels of OXT and OXT receptor (OXTR), following hsa-miR-193b overexpression, were analyzed by qRT-PCR (B and C) and western blot analysis (D). (E and F) Transcriptional and translational levels of MMP-2 and MMP-9, following hsa-miR-193b overexpression, were analyzed by qRT-PCR (E) and western blot analysis (F). (G) Following hsa-miR-193b overexpression, western blot analysis was performed to analyze the protein expression levels of total ERK1/2 and phosphorylated ERK1/2. The mRNA and protein expression levels were normalized to those of β-Actin. The data are expressed as the mean ± standard deviation (SD), *P ≤ 0.05, relative to the negative control group.

Furthermore, the ectopic expression of hsa-miR-193b significantly reduced the mRNA and protein levels of MMPs, which are genes that are related to trophoblast penetration regulated by OXT signaling, in JEG-3 cells (Figure 2E and F). The overexpression of hsa-miR-193b also inhibited ERK1/2 phosphorylation, an upstream mechanism involved in the expression of MMPs (Figure 2G). In addition, we observed that hsa-miR-193b regulated various angiogenesis-related genes, including vascular endothelial growth factor (VEGF) and fms-like tyrosine kinase 1 (sFlt-1), in JEG-3 cells. Specifically, hsa-miR-193b downregulated the protein expression of VEGF while upregulating that of sFlt-1 (Figure S1B), which aligns with the clinical characteristics observed in the placenta of patients with PE. These findings suggest that the OXT signaling pathway can be inhibited by placenta-derived miRNA (miR-193b-5p) associated with PE, which leads to the dysregulation of placental invasion.

### Hypoxic conditions decrease trophoblastic invasion activity and vasculature formation by inhibiting OXT signaling

PE has been reported to be caused by inadequate oxidative stress in the placenta ^28^. Considering these previously described results, we further investigated the effect of hypoxia, a significant factor that contributes to PE, on the placenta OXT signaling. Hypoxia was induced in JEG-3 cells to mimic the hypoxic conditions associated with PE, and the impact of OXT on the placental cells was evaluated.

Upon the induction of hypoxia in JEG-3 cells, the OXT concentration in the cell culture medium and the mRNA expression levels of OXT were found to be significantly decreased (Figure 3A and B), whereas there was no change in the expression level of OXTR (Figure S2A). To assess the effect of hypoxia on invasion activity, which is closely correlated with OXT signaling in JEG-3 cells, we examined the protein expression of HIF1A, a well-known marker for the detection of hypoxia ^29^. As expected, the HIF1A protein levels were markedly elevated in the hypoxic group (Figure 3C). However, pretreatment with OXT downregulated this increase in HIF1A protein expression (Figure 3D), and even under normoxic conditions, OXT tended to decrease the HIF1A levels (Figure S2B). Hypoxia induction downregulated the protein expression of OXTR and MMPs in JEG-3 cells, whereas the MMP levels were restored in the OXT pretreatment group (Figure 3E and 3F). In addition, we analyzed the effect of hypoxia on miR-193b-5p expression, which is significantly associated with OXT signaling in JEG-3 cells. As expected, the hsa-miR-193b-5p expression levels in JEG-3 cells under hypoxic conditions were significantly increased compared with those of the control group (Figure 3G).

**Figure 3.**
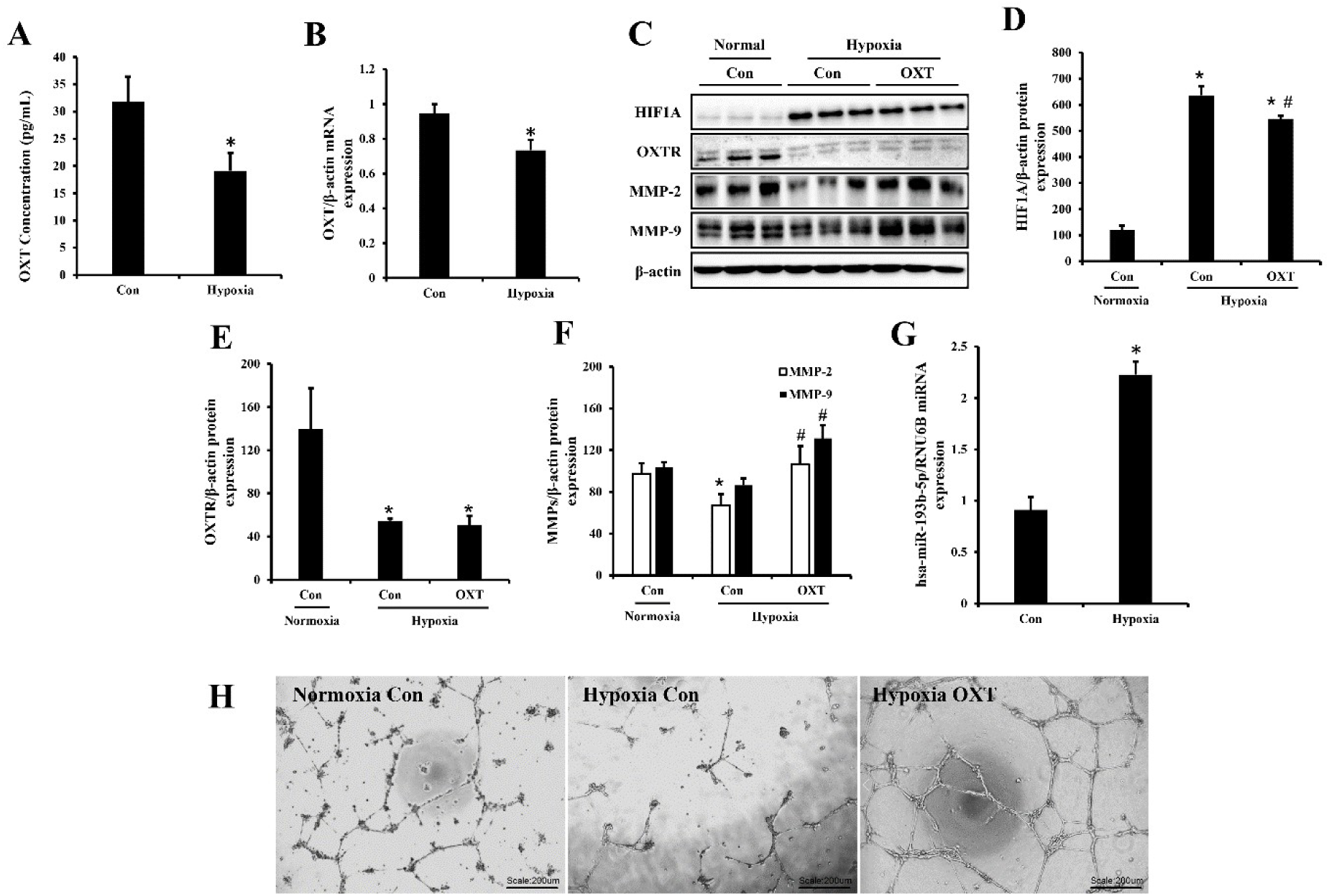
Inhibition of oxytocin (OXT) signaling by hypoxia reduces trophoblastic permeability and vascularization. (A) OXT concentration (pg/mL) in the cultured medium of JEG-3 cells under hypoxic conditions was analyzed by an ELISA assay. The data are expressed as the mean ± standard deviation (SD), *P ≤ 0.05, relative to the control group. (B) Gene expression analysis was performed using OXT-specific primers. β-Actin was used to normalize the gene expression. The data are expressed as the mean ± SD, *P ≤ 0.05, relative to the control group. (C) The JEG-3 cells were cultured in the presence or absence of OXT under hypoxic conditions. A western blot was performed to analyze the protein expression levels of HIF1A, OXTR, MMP-2, and MMP-9. Representative blot images are shown. (D-F) The protein expression levels of HIF1A (D), OXT receptor (OXTR) (E), and matrix metalloproteinases (MMPs) (MMP-2 and MMP-9) (F) were quantified and normalized to those of β-Actin. The data are expressed as the mean ± SD. *P ≤ 0.05 compared to the normoxia control group; ^#^P ≤ 0.05 compared to the hypoxia control group. (G) (C) The expression levels of hsa-miR-193b-5p in hypoxia-induced JEG-3 cells were determined by qRT-PCR. The small nuclear RNA RNU6B served as an internal control. The data are expressed as the mean ± SD, *P ≤ 0.05, relative to the control group. (H) HUVECs were seeded onto polymerized Matrigel as described in the Materials and Methods Section. The cells were cultured in the presence or absence of OXT under hypoxic conditions. Changes in cell morphology were captured through a microscope. Representative images show endothelial cell tube formation. Scale bar = 200 µm.

Placental angiogenesis is crucial for placental development and a successful pregnancy^30^. The impairment of placental angiogenesis, which is caused by oxidative stress, is involved in inadequate implantation and various pathological symptoms, including maternal anemia, placental growth restriction, and PE ^31^. Considering these aforementioned results, we examined the effect of OXT on hypoxia-induced endothelial dysfunction, a pathogenic factor in PE, by evaluating the endothelial tube formation in human umbilical vein endothelial cells (HUVECs). This method is widely used as an *in vitro* assay for assessing angiogenesis ^32–34^. As depicted in Figure 3G, the tube-like structures formed by HUVECs that were cultured under hypoxic conditions exhibited a reduced vasculature formation capacity compared with those cultured under normoxic conditions. However, HUVECs exposed to hypoxia and pretreated with OXT exhibited a recovery in tube formation compared with the non-treated hypoxic group (Figure 3H). Previous studies have reported that OXT induces tube formation in HUVECs ^35,36^, and as demonstrated in Figure S2C, our results also revealed that OXT increased tube formation in HUVECs under normoxic conditions. Collectively, these findings indicate that hypoxic conditions negatively impact OXT signaling, which results in the downregulation of placental invasive activity and angiogenesis.

### Inhibition of OXT signaling exhibits signs and symptoms mimicking PE in pregnant rats

The pregnant rats were administered the OXTR antagonist atosiban at doses of 0.6 mg/kg/day and 1.2 mg/kg/day from gestational day (GD) 10 to 17. To evaluate the physiological condition of the rats, maternal body weight (BW), placenta tissues, and fetuses were measured on GD 18 (Table S4). The maternal BW and fetal weight were significantly reduced in the atosiban 1.2 mg/kg/day group compared with the control group. Furthermore, the representative indicators of PE, including blood pressure levels, proteinuria concentrations, and renal pathology, were analyzed to determine whether the inhibition of OXT signaling during pregnancy induced PE characteristics and symptoms.

The blood pressure measurements indicated that the systolic and diastolic blood pressures in all the experimental groups remained within the physiologically normal range of approximately 120 mmHg and 90 mmHg, respectively ^37^, with no significant changes observed until GD 18 (Figure 4A). However, urine protein excretion was significantly higher in the atosiban 1.2 mg/kg/day group than in the control group (Figure 4B). Histological analysis of the kidneys revealed pathological changes in the renal tissues, including glomerular enlargement with occlusion of the capillary loops by swelling and hypertrophy of the endocapillary cells, in the 1.2 mg/kg/day atosiban group (Figure 4C). These findings demonstrate the pregnancy-specific dysfunction of the kidney, which was characterized by the extensive capillary occlusion referred to as “glomerular endotheliosis” ^38^.

**Figure 4.**
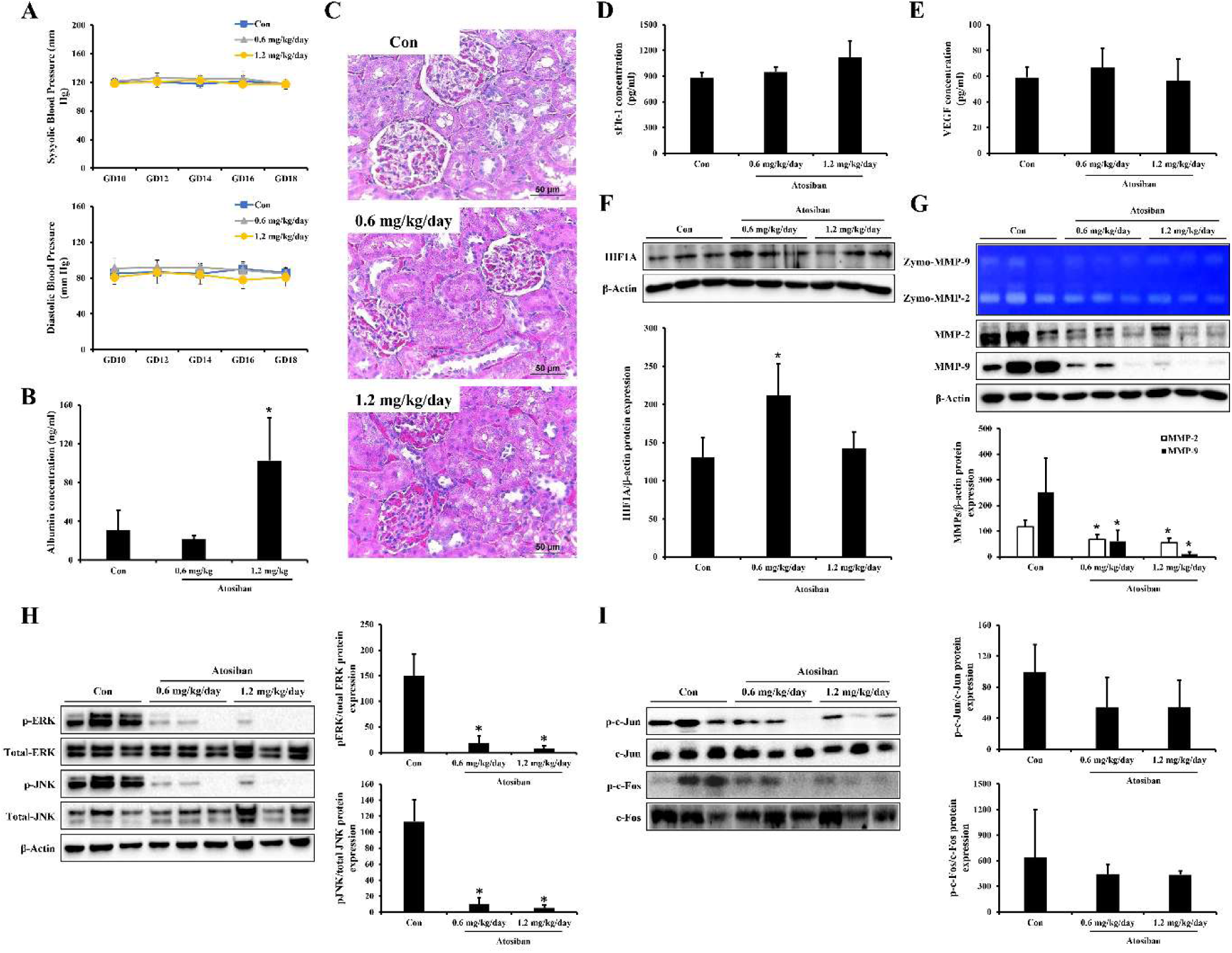
Atosiban inhibits the placental invasion mechanism in pregnant animals and induces signs and symptoms that mimic preeclampsia (PE). (A) The systolic and diastolic blood pressure levels in the atosiban-administered groups were measured and compared to the control group. The baseline blood pressures were obtained using a blood pressure monitor after the animals were pre-warmed on a warming platform in rat holders. Blood pressure readings were measured in triplicate, and the mean for each rat was recorded until gestation day (GD) 18. (B) Urine albumin concentration was analyzed using a rat albumin ELISA kit. Proteinuria was significantly observed in pregnant rats administered 1.2 mg/kg/day of atosiban. The data are expressed as the mean ± standard deviation (SD), *P ≤ 0.05, relative to the control group. (C) The kidneys were stained with hematoxylin & eosin. Staining results show capillary occlusion in pregnant rats administered with 1.2 mg/kg/day of atosiban with enlarged glomeruli and swollen endothelial cells compared to the control group. Scale bar = 50 μm. (D and E) Serum concentrations of sFlt-1 (D) and VEGF (E) in pregnant rats administered with atosiban were analyzed by ELISA and compared to the control group. (F) The expression level of placental HIF1A, a potential biomarker for predicting the risk of PE, was analyzed by western blot analysis, and the values are represented as schematic graphs. The individual protein expression levels were normalized to those of β-Actin. The data are expressed as the mean ± SD, *P ≤ 0.05, relative to the control group. (G) The activities and protein expression levels of the matrix metalloproteinases (MMPs) (MMP-2 and MMP-9) in the placenta of rats administered with atosiban were analyzed by gelatin zymography and western blot analysis, and the values are represented as schematic graphs. The individual protein expression levels were normalized to those of β-Actin. The data are expressed as the mean ± SD, *P ≤ 0.05, relative to the control group. (H) The protein expression levels of phosphorylated ERK1/2/total ERK1/2 and phosphorylated JNK/total JNK in the placenta of the pregnant rats administered with atosiban were quantified, and the values are represented as schematic graphs. The data are expressed as the mean ± SD, *P ≤ 0.05, relative to the control group. (I) The protein expression levels of p-c-Jun/c-Jun and p-c-Fos/c-Fos were analyzed by western blot analysis, and the values are represented as schematic graphs.

Next, we analyzed the sFlt-1 and VEGF serum levels, which are known biomarkers of PE. The administration of atosiban from GD 10 to 17 did not significantly change the serum levels of sFlt-1 and VEGF in the pregnant rats, although a slight increase in sFlt-1 levels was observed (Figure 4D and E). The protein expression levels of HIF1A, a potential predictive biomarker of PE associated with abnormal oxidative stress in the placental tissue, were increased by the atosiban 0.6 mg/kg/day group but not in the atosiban 1.2 mg/kg/day group compared with the control group (Figure 4F).

### Atosiban inhibits the placental invasion mechanism by blocking the ERK1/2 and JNK signaling pathways

To evaluate the impact of atosiban, an OXT signaling inhibitor, on placental function, we investigated the MMP activity and protein expression levels in the placenta of rats that were treated with atosiban. The gelatin zymography assay demonstrated that atosiban inhibited MMP activity in a dose-dependent manner, as shown in Figure 4G. Furthermore, the protein expression levels of placental MMPs were significantly decreased in all atosiban-treated groups compared with the control group (Figure 4G).

Based on our *in vitro* findings (Figure 1D and E), we hypothesized that atosiban could suppress MMP expression by inhibiting the ERK1/2 and JNK pathways in the placenta. To validate this hypothesis, we examined the effects of atosiban on ERK1/2 and JNK phosphorylation, which were significantly reduced in a dose-dependent manner (Figure 4H). In addition, we discovered that the Akt/mTOR pathway, another intracellular signaling pathway involved in various biological and pathological processes, including invasion activity ^39,40^, was also inhibited by atosiban (Figure S3). Finally, we assessed the inhibitory effect of atosiban on the activation of c-Jun and c-Fos, downstream components of the ERK1/2 and JNK pathways, and key regulators of MMP transcription. As depicted in Figure 4I, c-Jun and c-Fos phosphorylation tended to decrease in a dose-dependent manner in the placenta of rats treated with atosiban.

Collectively, these findings suggest that atosiban administration during pregnancy resulted in the inhibition of OXT signaling in the placenta and suppressed the ERK1/2/AP-1 and JNK/AP-1 pathways, which ultimately resulted in reduced invasion activity of the placenta by reducing both the expression and activity of MMPs.

### 2.1 OXT signaling was inhibited in an in vivo rat model of PE

Various *in vitro* and *in vivo* models have been developed to investigate the various aspects of PE ^41–43^. The reduction in uterine blood flow because of abnormal trophoblast invasion of the spiral arteries is a known cause of PE, and the reduced uterine perfusion pressure (RUPP) model is a suitable animal model for studying PE ^44^. The RUPP model induces several physiological features of PE, including hypertension and proteinuria. It also increases the plasma levels of sFlt-1 and decreases the plasma levels of VEGF and PlGF ^45,46^.

In this study, we utilized pregnant rats to establish a RUPP model and investigated the effects of abnormal placental hypoxic stress on OXT signaling during pregnancy. Immunofluorescence analysis was performed on the placenta tissue using antibodies specific against OXT and OXTR to evaluate the changes in OXT signaling caused by RUPP. As shown in Figure 5A, the RUPP placentas exhibited decreased staining for OXT and OXTR compared with the SHAM placenta tissues. Interestingly, the immunofluorescent localization of the target proteins, including OXT, was most prominent in the placental labyrinth compared with the decidua and junctional zone (Figure S4A–D). The placental labyrinth serves as an interface for gaseous and nutrient exchange between the embryo and the mother ^47,48^.

**Figure 5.**
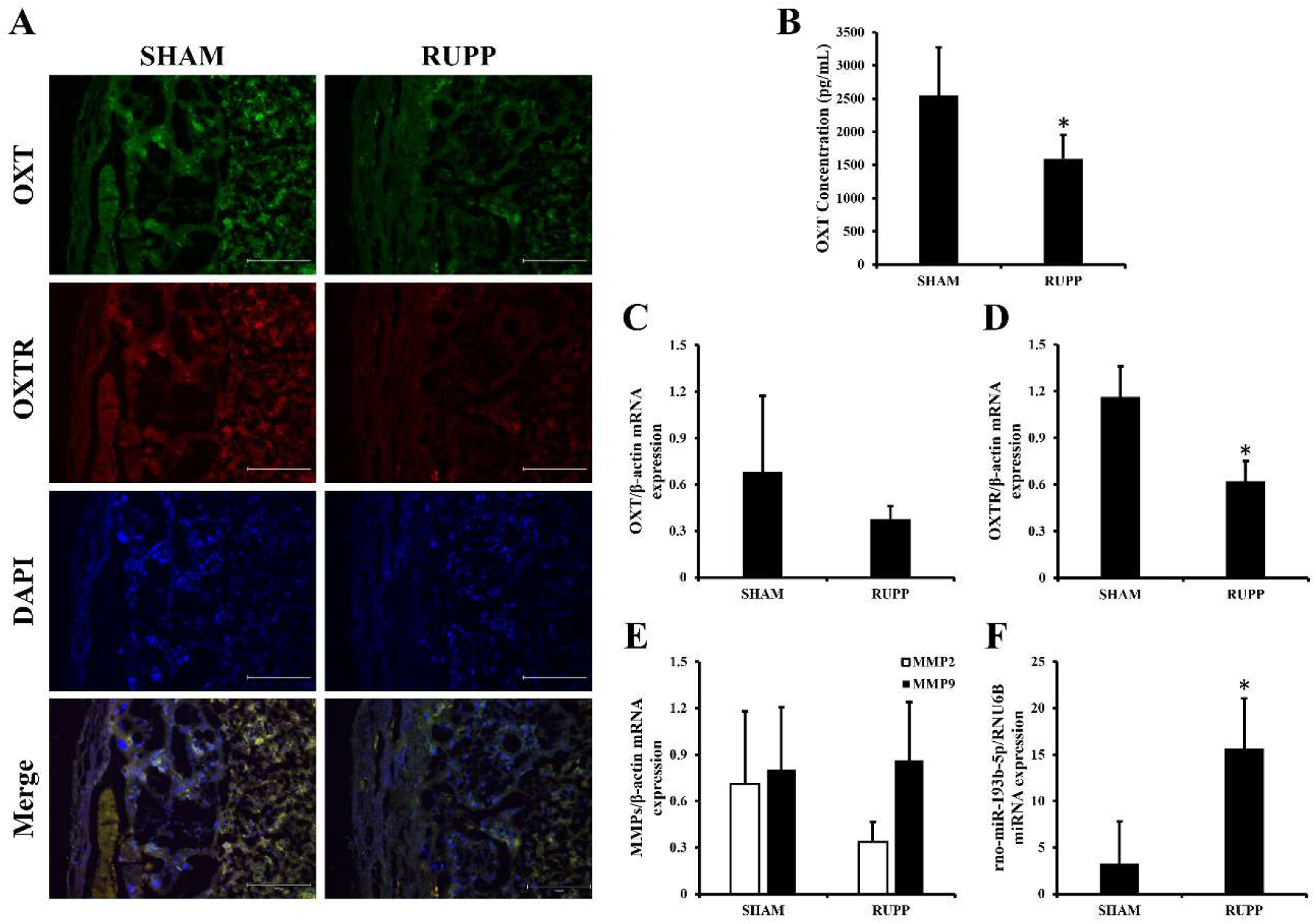
The oxytocin (OXT) signaling pathway was attenuated in the reduced uterine perfusion pressure (RUPP) rat model. (A) Pregnant rats underwent a RUPP operation to establish a preeclampsia (PE) *in vivo* model. At the end of the experiment, the harvested and fixed placenta tissues from the rats were subjected to immunofluorescent staining targeting OXT (green) and OXT receptor (OXTR) (red). Nuclei were stained with DAPI (blue). Representative images show the abnormal expression of OXT and OXTR in the placenta of the RUPP rat model. Scale bar = 300 µm. (B) The serum OXT concentration of pregnant rats that underwent a RUPP operation was analyzed by ELISA and compared to the control group. *P ≤ 0.05 compared to the SHAM group. (C-E) Total mRNA was extracted from the placenta of pregnant rats that underwent a RUPP operation. The placental mRNA expression levels of OXT (C), OXTR (D), and matrix metalloproteinases (MMPs) (MMP-2 and MMP-9) (E) were analyzed by qRT-PCR. β-Actin was used to normalize the gene expression. *P ≤ 0.05 compared to the SHAM group. (F) The expression level of rno-miR-193b-5p in the placenta of the RUPP rat model was determined by qRT-PCR. The small nuclear RNA RNU6B served as an internal control. The data are expressed as the mean ± standard deviation (SD), *P ≤ 0.05 compared to the SHAM group.

Furthermore, the RUPP group exhibited a significantly lower blood OXT concentration than the SHAM group (Figure 5B), and there was a relative reduction in the mRNA expression of OXT (Figure 5C). The mRNA expression of OXTR was also significantly decreased in the placenta of the RUPP group (Figure 5). However, the MMP mRNA expression levels did not change significantly (Figure 5E). We further analyzed the involvement of miRNA-193b-5p in OXT regulation in the RUPP model. Figure 5F demonstrates that the expression of rno-miR-193b-5p was significantly increased in the placenta of the RUPP group compared with the SHAM group.

Considered together, these findings suggest that the induction of abnormal placental hypoxia through the RUPP model inhibits the OXT signaling pathway, which is consistent with the *in vitro* results. This increased miR-193b-5p, a major indicator of PE in the placenta.

### Placental implantation abnormalities induced by an OXT signaling inhibitor in pregnant rats

Implantation, the process by which placental cells penetrate the uterine wall during the early stages of pregnancy, is crucial for maintaining pregnancy ^49^. Previous studies have indicated the importance of OXT throughout pregnancy ^18^, and we hypothesized that OXT was essential during implantation. In this study, we administered subcutaneous injections of atosiban (1.2 mg/kg/day) to pregnant rats from GD 3 to 7 to evaluate the effect of OXT signaling inhibition during the implantation period. At the end of the experiment, we assessed the influence of atosiban on the implantation rate by examining the number of implantation sites in the uterus. However, we did not observe any significant changes in the number of implantation sites upon OXT signaling inhibition (Table S5).

Next, we investigated the effects of atosiban on the protein expression levels of the MMPs and HoxA-10, which are genes that are crucial for placental transplantation during implantation. Atosiban administration downregulated the MMP-9 and HoxA-10 protein expression levels in the uterus but not MMP-2 (Figure S6A–C). Furthermore, we analyzed the VEGF and sFlt-1 protein expression levels, which are associated with angiogenesis and serve as major biomarkers of PE. The administration of atosiban during early pregnancy significantly reduced the VEGF protein levels, whereas the sFlt-1 protein levels exhibited no significant change (Figure S6D–F).

These findings suggest that OXT also plays an important role in placental implantation during the early stages of pregnancy and may be a valuable indicator for the early diagnosis of PE.

### OXT is a potential biomarker for the early diagnosis of PE

Considering the results described above, we proceeded to investigate the correlation between patients with PE and OXT signaling. We analyzed the mRNA and protein expression levels of OXT and OXTR in placenta tissue obtained from patients with PE. Our results demonstrated a significant decrease in the mRNA and protein expression levels of OXT and OXTR in the PE group compared with the normal group (Figure 6A and B). Furthermore, we examined the plasma levels of endogenous OXT in patients with PE using an ELISA assay. The plasma OXT concentration in patients with PE during their third trimester (29–40 weeks) of pregnancy was found to be 3.5 times higher than that of the normal group (Figure 6C).

**Figure 6.**
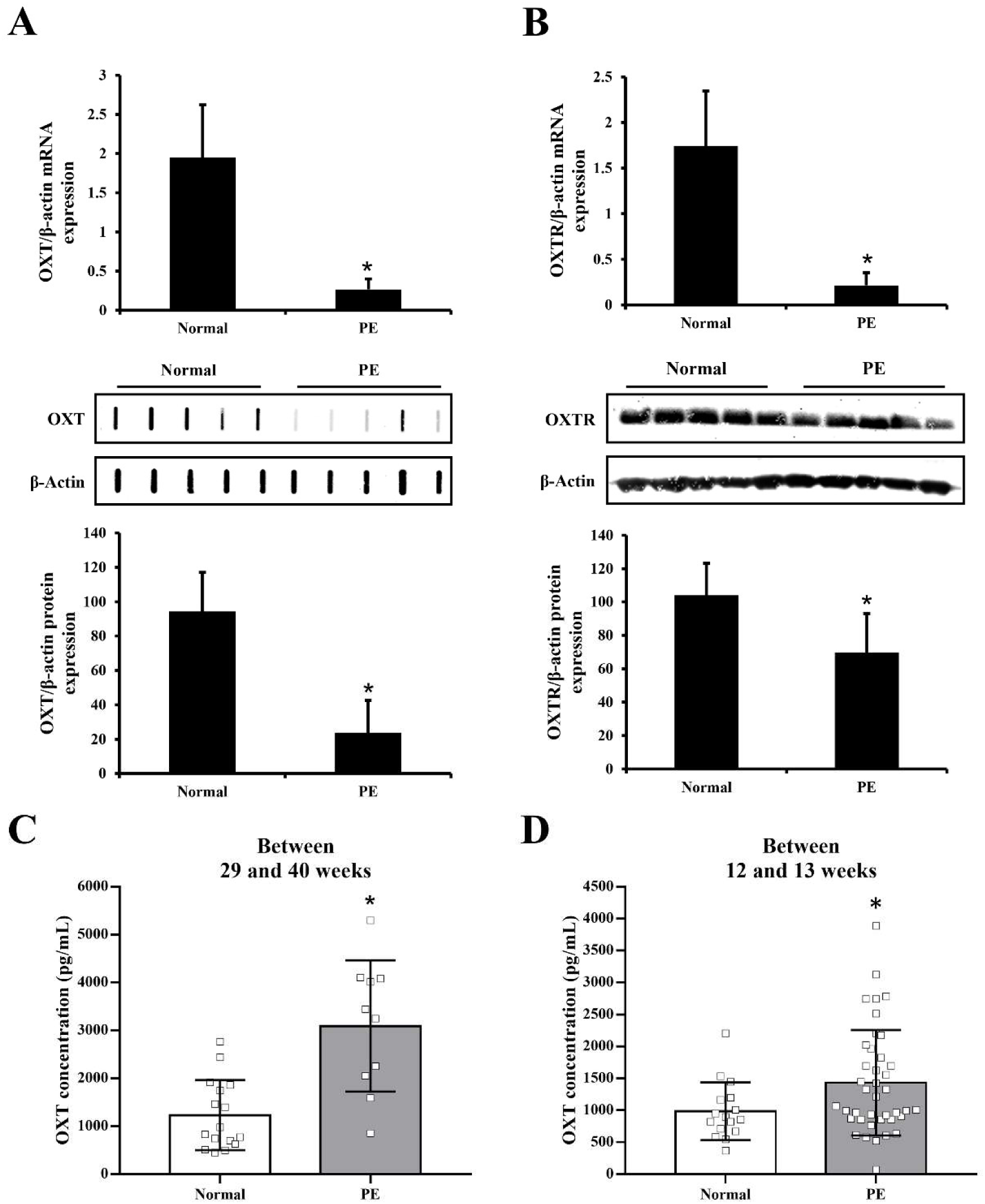
Oxytocin (OXT) signaling is significantly regulated throughout pregnancy in patients with PE. (A and B) The total mRNA and proteins extracted from whole placental tissues during late pregnancy (between 29 and 40 weeks) of normal women (n = 21) and patients with PE (n = 20) were analyzed for expression of OXT (A) and OXT receptor (OXTR) (B) using qRT-PCR, dot-blot analysis, and western blot analysis. β-Actin was used to normalize the gene expression. *P ≤ 0.05 compared to the normal women group. (C and D) Plasma OXT concentration (pg/mL) in late pregnancy (between 29 and 40 weeks) (C) and early pregnancy (between 12 and 13 weeks) (D) of patients with PE were analyzed by ELISA assay. The data are expressed as the mean ± standard deviation (SD). *P ≤ 0.05 compared to the normal women group.

To propose OXT as a diagnostic marker for PE during early pregnancy, we collected blood samples from pregnant women during their first trimester (12–13 weeks). Among the pregnant women who provided blood samples, only 40 individuals exhibited PE symptoms in the later stages of pregnancy (after 20 weeks) ^50^. The clinical characteristics of the patients with PE are detailed in Table S2. Interestingly, the plasma OXT concentrations in patients with PE during their first trimester were significantly higher than those of the normal group (Figure 6D). These results strongly suggest that OXT signaling was significantly inhibited in the placenta of patients with PE. Moreover, we demonstrated that OXT holds great potential as a novel biomarker for the early diagnosis of PE during the first trimester of pregnancy.

## Discussion

Considerable research efforts have been dedicated to understanding PE, a condition whose treatment and causes remain elusive. A recent report recommended the use of aspirin as a preventive therapy for PE, while statins are being investigated as potential treatments ^51^. Notably, the G-protein-coupled receptor (GPCR) family has emerged as a vital contributor to maternal physiological adaptations during pregnancy and placental development. Dysregulated GPCR signaling can affect the progression of severe PE ^52^. Several GPCR targets show promise in improving PE symptoms and safely extending the pregnancy, which is the ultimate goal of PE treatment ^53^. Among these, OXTR, a member of the rhodopsin-type GPCR family, is activated through its interaction with OXT ^54^. The OXT-OXTR system plays a pivotal role throughout pregnancy. Our previous study confirmed that OXT signaling levels in the placenta gradually increase during the late stages of pregnancy ^18^. However, despite reports of OXT signaling synthesis and expression in the placenta, its precise role therein remains incompletely understood.

Several lines of evidence strongly suggest that OXT signaling is intricately linked to placental function and that its dysfunction can substantially influence the development of PE symptoms. In this study, we demonstrated the effect of OXT on activating the MMP signaling pathway in placenta cells, which is critical for successive placental invasion into uterus. The regulation of MMP by OXT was stimulated through the increased phosphorylation levels of the c-Jun, c-Fos, ERK1/2, and JNK pathways. The ERK1/2 and JNK pathways activated AP-1, the critical transcription factor of MMPs. Additionally, we established that inhibiting OXT signaling markedly reduced placental permeability, which was attributed to the inhibition of the MMP signaling pathways. Disrupting OXT signaling by the administration of the OXTR inhibitor atosiban to pregnant animals induces preeclamptic symptoms such as placental hypoxia, proteinuria, and renal impairment. Notably, a previous study demonstrated that MMP-9 silencing in knockout mice exhibited a phenotype resembling PE, likely because of impaired trophoblast differentiation and invasion activity ^55^. In addition, downregulated expression levels of MMPs have been observed in preeclamptic placentas compared with normal placentas^56,57^.

To confirm the role of OXT during implantation, we treated pregnant rats with atosiban during the implantation period of pregnancy. Atosiban decreased the expression of the implantation-associated proteins, including MMP-9, HoxA-10, and VEGF in the uterus, which suggests that deficiencies in OXT signaling adversely influence implantation and placental development. The cause of PE is not clearly suggested. However, inappropriate implantation during early pregnancy causes a lack of blood flow to the fetus from the mother, which increases the blood pressure in the mother to supply more blood to the fetus. Indeed, the symptoms of PE, such as high blood pressure, maternal renal impairment, and placental hypoxia, are associated with the effort of the mother to increase blood flow to the fetus ^58^. Our findings highlight that alteration in OXT signaling during the early stage of pregnancy can lead to disturbances in implantation, which induce abnormal placental growth and function, resulting in the development of PE during the later stages of pregnancy.

Various factors influence the development of PE. Using bioinformatic prediction programs, we initially predicted that miR-193b-5p could influence OXT gene expression. This miRNA that is derived from the placentas of patients with PE is among those identified in recent studies as potential contributors to the onset of PE ^59,60^. We confirmed that miR-193b-5p suppressed OXT expression in JEG-3 cells and inhibited the expression of key genes, such as MMP-2 and MMP-9, which play crucial roles in placental-uterine invasion. Placental hypoxia has been linked to reduced placental invasion capacity and endothelial cell dysfunction and significantly contributes to the pathogenesis of PE ^61–63^. Our findings revealed that OXT mitigates the reduction of placental invasion capacity and angiogenesis caused by hypoxia. Several studies have also reported that OXT stimulates the migration and invasion of HUVECs through the activation of OXTR, which is expressed in the endothelial cells ^64,65^. Therefore, we examined the OXT signaling pathway in a RUPP model, which is a representative PE-mimicking animal model by inducing hypoxia in the placenta. Intriguingly, the hypoxic conditions in the placenta significantly elevated the expression of miR-193b-5p and OXT signaling. These results indicate that OXT is a critical endocrine factor associated with PE. However, it is important to note that although the RUPP model shares some characteristics with PE, it has limitations in fully representing the complexity of human PE because the rat placenta is anatomically and physiologically different from the human placenta ^66,67^.

Our experimental findings, gathered from *in vitro* and *in vivo* data, establish a clear and significant correlation between OXT signaling and PE. Motivated by these results, we extended our investigation to assess the relevance of OXT signaling in patients with PE. We verified that the expression levels of OXT and its receptor, OXTR, were reduced in the placentas of patients with PE. These findings support the concept that the dysfunction of OXT signaling plays a vital role in the onset of the disease by compromising placental functions. Furthermore, we observed a significant increase in plasma OXT levels during the later stages of pregnancy in the PE group. This observation aligns with a previous study, which indicated that plasma OXT levels in full-term pregnancy increase with hypertension severity ^68^. To explore the possibility of using OXT as a diagnostic marker during the early stages of pregnancy, we assessed the plasma OXT levels in blood samples collected from women during their first trimester who later developed PE during late pregnancy. Our data suggests that pregnant women with elevated levels of plasma OXT during early pregnancy have a greater potential to develop PE during late pregnancy. These findings collectively emphasize the potential use of OXT as a diagnostic marker for the early identification of PE, thereby emphasizing its relevance in clinical practice.

### Perspective

Oxytocin (OXT), traditionally known for its role in parturition and lactation, is increasingly recognized as a regulator of placental development. This study presents compelling evidence that OXT signaling is essential for proper trophoblast invasion and vascular development via activation of MMP-2 and MMP-9 through ERK1/2 and JNK/AP-1 pathways. Disruption of OXT signaling, whether by miR-193b-5p, hypoxic conditions, or pharmacological inhibition, impairs placental function and induces preeclampsia (PE)-like features in vivo. The demonstration that maternal OXT levels are elevated during the first trimester in women who later develop PE suggests a compensatory endocrine response to early placental dysfunction, positioning OXT as a promising early biomarker for PE prediction. These findings provide mechanistic insight into the hormonal regulation of placental invasion and angiogenesis and support the idea that PE originates from impaired implantation and trophoblast differentiation. Importantly, the involvement of the RUPP and atosiban-treated animal models enhances the translational relevance of the data. Future research should aim to validate these findings in larger clinical cohorts, assess the predictive power of OXT in combination with other biomarkers, and explore therapeutic strategies that target the OXT-OXTR axis to restore placental function. If successful, such strategies could enable earlier diagnosis and more effective management of PE, ultimately improving outcomes for both mothers and their babies. This study broadens our understanding of pregnancy-related hypertensive disorders and emphasizes the importance of placental hormone signaling in maternal-fetal health.

### Novelty and Relevance

#### What Is New?

This study identifies oxytocin (OXT) as a key regulator of trophoblast invasion, acting through the ERK1/2 and JNK/AP-1 pathways to enhance MMP-2 and MMP-9 expression. We demonstrate that hsa-miR-193b-5p, a placenta-derived microRNA associated with preeclampsia (PE), suppresses OXT signaling and impairs invasion. Notably, we show for the first time that maternal plasma OXT levels are significantly elevated during the first trimester in women who later develop PE. This suggests a potential compensatory mechanism and supports the use of OXT as a novel biomarker for early prediction of PE before clinical symptoms arise, providing new mechanistic insight.

#### What Is Relevant?

Our findings directly link oxytocin signaling to mechanisms underlying hypertensive disorders of pregnancy, particularly preeclampsia. Impaired OXT signaling, caused by hypoxia or miR-193b-5p, disrupts placental invasion and angiogenesis—hallmarks of PE pathogenesis. This study further validates the role of OXT using two in vivo models: a reduced uterine perfusion pressure (RUPP) model and an OXT receptor antagonist (atosiban) model, both recapitulating PE-like symptoms. Together, this evidence strengthens the connection between hormonal signaling dysfunction and hypertensive pregnancy complications, highlighting the clinical relevance of the OXT pathway in understanding and monitoring the development of PE.

#### Clinical/Pathophysiological Implications?

The discovery that oxytocin levels are elevated during the first trimester in women who later develop preeclampsia suggests its clinical utility as an early, non-invasive biomarker. By enabling early detection before symptom onset, OXT screening may allow for timely risk stratification and intervention, which are crucial for improving maternal and fetal outcomes. Furthermore, therapeutic strategies aimed at restoring or modulating OXT signaling could represent a novel approach to mitigating placental dysfunction. These findings offer new insights into the pathophysiological mechanisms of PE and present opportunities for more effective prediction, prevention, and treatment strategies in hypertensive pregnancy disorders.

#### Ethics approval and consent to participate

This study was approved by the Institutional Review Board (IRB) of the Pusan National University Hospital Clinical Trial Center (1302-005-015, 2006-030-092), and all participants provided written informed consent. Animal studies were approved (approval number: PNU-2022-0161) by the Ethics Committee of Pusan National University (Busan, Republic of Korea).

## Consent for publication

Not applicable.

## Availability of data and materials

This study includes no data deposited in external repositories. Availability of data and materials.

## Author contributions

Sung-Min An: Conceptualization, Methodology, Data curation, Formal analysis, Investigation, Visualization, Writing – original draft, Writing – review & editing. Jea Sic Jeong: Conceptualization, Methodology, Data curation, Formal analysis, Investigation, Visualization, Writing – original draft, Writing – review & editing. Min Jae Kim: Validation, Formal analysis, Investigation, Writing–review & editing. Da Som Kim: Validation, Formal analysis, Investigation, Writing – review & editing. So Young Kim: Validation, Formal analysis, Investigation. Hyeon-Gu Kang: Validation, Formal analysis, Investigation. Seung Chuel Kim: Conceptualization, Resources, Supervision, Methodology, Data curation, Formal analysis, Investigation, Writing – original draft, Writing – review & editing. Beum-Soo An: Conceptualization, Funding acquisition, Project administration, Resources, Supervision, Methodology, Data curation, Formal analysis, Investigation, Writing – original draft, Writing – review & editing.

## Acknowledgements

This work was supported by the National Research Foundation of Korea(NRF) grant funded by the Korea government(MSIT) (No. 2020R1F1A105243311) and partially supported by the BK21 FOUR Program(4120240915200) through the National Research Foundation of Korea (NRF), funded by the Ministry of Education, Korea

## Supplemental Material

Supplemental methods

Figure S1-S5

Table S1-S5

